# Breaking bud: probing the scalability limits of phylogenetic network inference methods

**DOI:** 10.1101/056572

**Authors:** Hussein A Hejase, Kevin J Liu

## Abstract

**Background:** Branching events in phylogenetic trees reflect strictly bifurcating and/or multifurcating speciation and splitting events. In the presence of gene flow, a phylogeny cannot be described by a tree but is instead a directed acyclic graph known as a phylogenetic network. Both phylogenetic trees and networks are typically reconstructed using computational analysis of multi-locus sequence data. The advent of high-throughput sequencing technologies has brought about two main scalability challenges:(1) dataset size in terms of the number of taxa and (2) the evolutionary divergence of the taxa in a study. The impact of both dimensions of scale on phylogenetic tree inference has been well characterized by recent studies; in contrast, the scalability limits of phylogenetic network inference methods are largely unknown. In this study, we quantify the performance of state-of-the-art phylogenetic network inference methods on large-scale datasets using empirical data sampled from natural mouse populations and synthetic data capturing a wide range of evolutionary scenarios.

**Results:** We find that, as in the case of phylogenetic tree inference, the performance of leading network inference methods is negatively impacted by both dimensions of dataset scale. In general, we found that topological accuracy degrades as the number of taxa increases; a similar effect was observed with increased sequence mutation rate. The most accurate methods were probabilistic inference methods which maximize either likelihood under coalescent-based models or pseudo-likelihood approximations to the model likelihood. Furthermore, probabilistic inference methods with optimization criteria which did not make use of gene tree root and/or branch length information performed best-a result that runs contrary to widely held assumptions in the literature. The improved accuracy obtained with probabilistic inference methods comes at a computational cost in terms of runtime and main memory usage, which quickly become prohibitive as dataset size grows past thirty taxa.

**Conclusions:** We conclude that the state of the art of phylogenetic network inference lags well behind the scope of current phylogenomic studies. New algorithmic development is critically needed to address this methodological gap.

## Background

In recent studies, gene flow-the process by which genetic material is exchanged between different populations and/or species existing at the same point in time-has been shown to have played a major role in the evolution of a diverse array of metazoans, including humans and ancient hominins [1, 2], mice [3], and butterflies[4]. Each of these organisms (as well as many others across the Tree of Life [5, 6, 7]) has a phylogeny, or evolutionary history, which cannot be represented as a tree, where a branching event reflects strict bifurcating and/or multifurcating speciation/splitting and subsequent genetic isolation of the resulting species/populations. In these cases, the phylogeny takes the more general form of a directed acyclic graph known as a phylogenetic network [8].

Similar to phylogenetic trees, phylogenetic networks are typically inferred using computational analyses of multi-locus biomolecular sequence data. The most widely used approach is a concatenated analysis which estimates a single phylogeny for all loci [9]. Methods used for this analysis typically only account for sequence mutation and gene flow [10]; all local genealogical discordance is ascribed to gene flow. Representative examples include NeighborNet [11] and the least squares method of Schliep [12], which we refer to here as SplitsNet. A primary complication with the concatenated approach is that different loci in a genome commonly exhibit local genealogical incongruence (i.e., gene trees can differ from each other and the species phylogeny in terms of topology and/or branch length) due to the complex interplay of different evolutionary processes that shaped the genomes. These include gene flow, sequence mutation, gene duplication and loss, recombination, and incomplete lineage sorting (ILS). ILS occurs when genetic drift causes lineages from two isolated populations to coalesce at a time more ancient than their most recent ancestral population, and is known to play a crucial role in the evolution of much of the Tree of Life [9].

In contrast to concatenated analysis, multi-locus methods infer species phylogenies in the presence of these evolutionary processes acting in combination. The most widely used multi-locus methods perform inference that account for a broad set of evolutionary processes, including sequence mutation, gene flow, and ILS [13, 14, 15]. Multi-locus methods are broadly classified by whether or not they impose the requirement that a phylogenetic hypothesis be specified *priori*.

The main focus of our study is the category of methods that perform full inference by searching among all possible phylogenies defined on a set of taxa. Many of these methods utilize a gene-tree/species-phylogeny reconciliation approach (or summary approach), where local trees estimated from different loci-referred to as gene trees-are used as input rather than sequence alignments from the loci [16, 17, 18, 19]. The full inference procedure therefore requires two phases: a first phase where a set of gene trees is estimated from biomolecular sequence alignments, and a second phase where the gene trees are used to estimate a species phylogeny. The multi-locus methods are further classified by the optimization criterion used for inference. Earlier parsimony-based approaches (e.g., the method of [20], which we refer to here as MP, which stands for maximum parsimony) utilize the minimize deep coalescence (MDC) criterion proposed by [8], which seeks the species phylogeny that minimizes the number of deep coalescences necessary to explain a given set of gene trees. More recently, probabilistic approaches perform phylogenetic network inference under an explicit evolutionary model that combines the coalescent model with biomolecular substitution models. Examples include two different methods proposed by [21] that are implemented in the PhyloNet software package [22], which differ primarily in their use of branch length information: one method uses the approach of [23] to calculate model likelihood using only gene tree topologies, and the other method substitutes an alternative approach to calculate model likelihood using gene tree topologies and branch lengths. We therefore refer to these methods as MLE (which stands for maximum likelihood estimation) and MLE-length, respectively. These probabilistic approaches have been noted to have high computational requirements, and model likelihood calculations were found to be a major performance bottleneck [24]. For this reason, pseudo-likelihood approximations to full model likelihood calculations have been proposed, including the method of [24] (referred to here as MPL, which stands for maximum pseudo-likelihood), which substitutes pseudo-likelihoods in the optimization criterion used by MLE, and SNaQ (Species Networks applying Quartets) [25], which combines the use of pseudo-likelihoods under a coalescent-based model with quartet-based concordance analysis [26]. As suggested by [14], the techniques used by [27] to infer a species tree directly from an input sequence alignment-effectively integrating over gene tree distributions at different loci-would provide an alternative to reconciliation-based species network inference, but scalable inference methods using this alternative approach have yet to be proposed and remain for future work as of this writing; preliminary experiments by [14] suggest that the scalability challenges of this approach will be greater than with state-of-the-art reconciliation-based approaches. All of these multi-locus methods address problems that are either known or suspected to be NP-hard [14, 15]. For this reason, heuristics are necessary to enable efficient inference under the optimization criteria associated with these methods. The practical design of the heuristics are essential to accuracy and computational efficiency.

A second category of methods requires a fixed phylogeny to be required as input. We note that the fixed-phylogeny inference problem is contained within the general phylogenetic inference problem. Fixed-phylogeny inference methods are typically used to address high-level questions such as detecting gene flow (e.g., the D-statistic [1] and its extensions [13]), inferring ancestral population sizes and other population genetic quantities (e.g., the CoalHMM method [28] which utilizes a hidden Markov model (HMM) to capture within-genome sequence dependence), and detecting genomic loci involved in gene flow (e.g., PhyloNet-HMM [29]). Since the primary focus of our study is the general phylogenetic network inference problem rather than special cases thereof, we do not consider these methods further.
Thanks to rapid advances in genome sequencing and related biotechnologies [30], large-scale phylogenetic studies involving many dozens of genomes or more are now common (see [31] for a survey). These developments pose two primary scalability challenges: (1) the number of taxa in a study, and (2) sequence divergence, which reflects the evolutionary divergence of the taxa in a study.

For the special case of phylogenetic tree inference from phylogenomic data, recent studies have examined these scalability challenges [32, 17, 33] (including evolutionary scenarios involving gene flow [34, 35]) and proposed new methods for large-scale analysis [32, 36, 37]. In contrast, for the more general case of phylogenetic network inference, the limits of scalability on inputs with more than a few dozen taxa as well as performance at these limits have yet to be established. What are the computational requirements of state-of-the-art methods, and what is their accuracy on large-scale inputs with dozens of taxa or more?

To resolve these open questions, we conducted a scalability study of state-of-the-art phylogenetic network inference methods on both simulated and empirical datasets. To our knowledge, our study is the first to address these open questions. We chose representative methods from the different categories discussed above: Neighbor-Net and SplitsNet (from the category of concatenated methods), MP (from the category of parsimony-based multi-locus inference methods), MLE and MLE-length (from the category of probabilistic multi-locus inference methods that use full likelihood calculations), and MPL and SNaQ (from the category of probabilistic multi-locus inference methods that use pseudo-likelihood approximations to the full model likelihood). Following the practice of prior simulation studies [25, 14], our performance comparison focuses on the simpler case of search among phylogenetic networks with the correct number of network nodes (which is one in all model conditions). The more general case of search among network hypotheses with differing network nodes necessitates the use of model selection techniques to balance model fit versus complexity, and is suspected to be more difficult for this reason [14, 38]. Our performance study utilized empirical data from past studies of natural mouse populations and synthetic data which were simulated to capture a wide range of evolutionary scenarios reflecting prior empirical studies. The performance of the phylogenetic network inference methods on the empirical and synthetic data was evaluated using three performance measures: (1) computational time, (2) memory usage, and (3) topological accuracy.

## Results

### Performance evaluation on simulated datasets

*Runtime and memory usage*. We began by assessing the effect of dataset size on computational time and memory requirements. We focused on the probabilistic multi-locus methods since they were the most accurate methods in our study (see below). Of the full likelihood methods in this category, we focused on MLE instead of MLE-length since the former is more accurate than the latter when inferred gene trees were used as input. The other methods-SNaQ and MPL-consisted of pseudo-likelihood-based approaches.

Based on sampled dataset sizes, the full likelihood method had runtime and memory usage that was strictly greater than the pseudo-likelihood-based methods (Figure 1i). The performance difference approached an order of magnitude. For all methods, runtime grew super-linearly as dataset size increased. The observed growth in runtime is consistent with previous performance studies [39, 40, 25]. For all methods, runtime became impractical on datasets that were much smaller than those used in current genomic studies. Given a maximum runtime of one week, the full likelihood and pseudo-likelihood-based methods were able to analyze datasets with 13 and 20 taxa, respectively. The two classes of methods required more than a week of runtime on datasets with 17 and 25 taxa, respectively. We also attempted analyses of datasets with 30, 40, 50, and 100 taxa; none of these analyses had finished after multiple weeks of runtime as of this writing.

**Figure 1.**
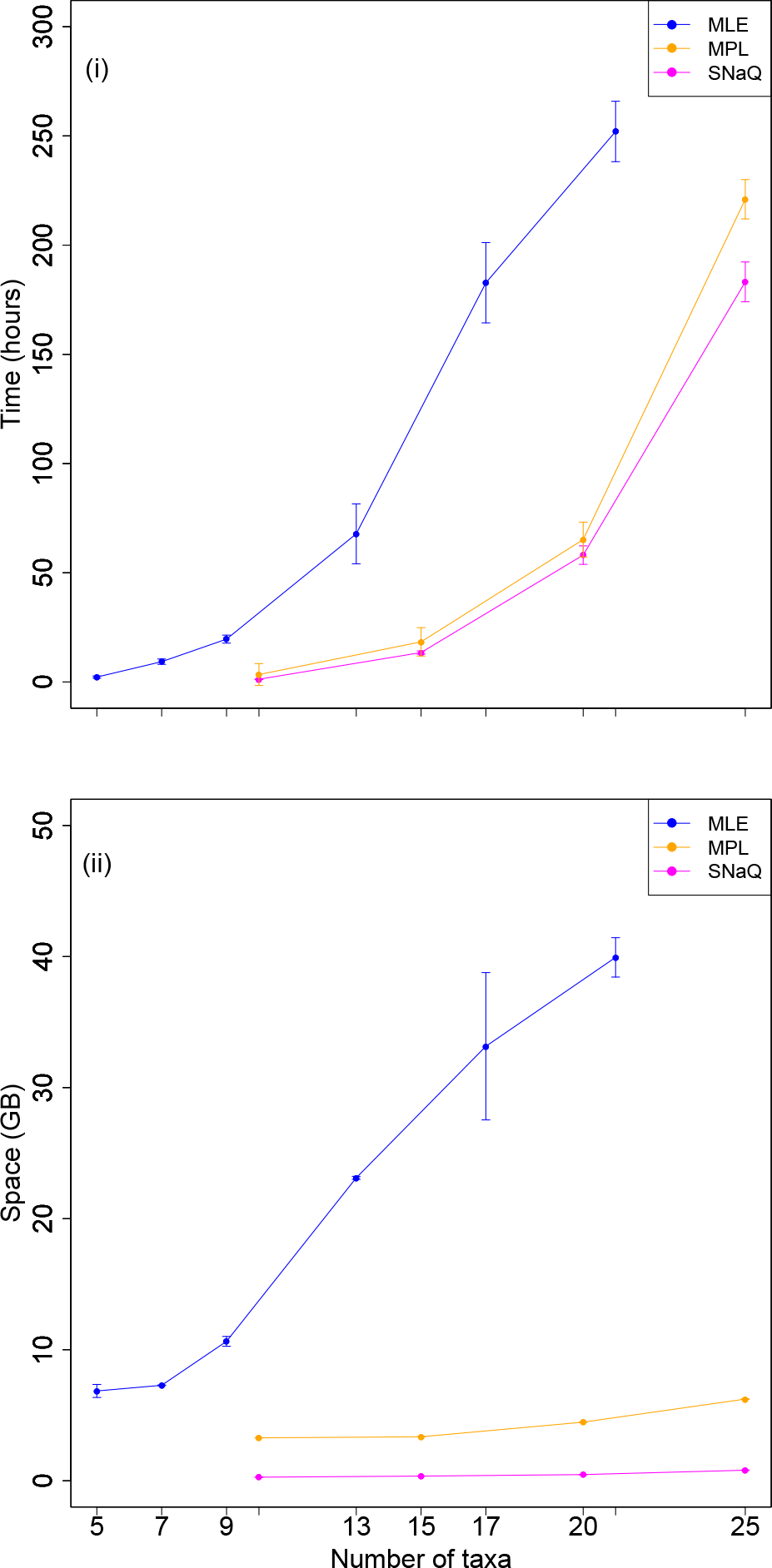
The impact of dataset size on the computational requirements of the probabilistic inference methods. Results are shown for MLE, MPL, and SNaQ analyses of simulated datasets. (i) Average runtime (h) and (ii) main memory usage (GiB) are shown with standard error bars (*n* = 5).

Relative to runtime performance, the main memory requirements of the different methods contrasted to a greater degree. On datasets with more than a dozen taxa, the full likelihood method exhibited super-linear growth in main memory usage, similar to its performance in terms of runtime (panel (ii) in Figure 1). MLE’s main memory requirements are projected to become prohibitive on datasets with more than a few dozen taxa. In contrast, the pseudo-likelihood-based methods had peak memory usage below 10 GiB on datasets with up to 25 taxa. MPL’s memory usage was flat as dataset size increased from 10 to 15 taxa, and increased by just a few GiB as dataset size increased from 15 to 25 taxa. SNaQ’s memory usage was largely constant at around a few GiB on datasets with sizes up to 25 taxa.

*Topological accuracy*.We next examined topological accuracy of the inference methods as dataset scale grew in two ways: the number of taxa and sequence divergence. We evaluated the topological accuracy of inferred phylogenetic networks using bipartition-based measures (see Methods) that generalize the Robinson-Foulds distance, a widely used bipartition-based distance defined on phylogenetic trees. We also explored an alternative tripartition-based measure [41]. Consistent with a prior study [25], we found that the tripartition-based measure was too sensitive for useful performance comparison: small topological edits tend to result in a large change in the tripartition-based distance. For this reason, we focus on comparisons using the bipartition-based measures.

As shown in Figures 2, 3 and 4, the methods fell into three categories based upon their topological accuracy: on all model conditions except for the model condition with the highest sequence divergence (the seven-taxon model condition with mutation rate θ = 1.6), (1) SNaQ was the most accurate, (2) MLE, MPL, and MP had intermediate accuracy relative to the other methods, and (3) MLE-length and the concatenated methods (Neighbor-Net and SplitsNet) were the least accurate methods. Note that, for each replicate, the same set of gene trees was provided to each multi-locus method as input. For each method, the largest topological error was seen on the largest datasets (relative to smaller datasets) and on the model conditions with the highest mutation rate (relative to model conditions with smaller mutation rates).

**Figure 2.**
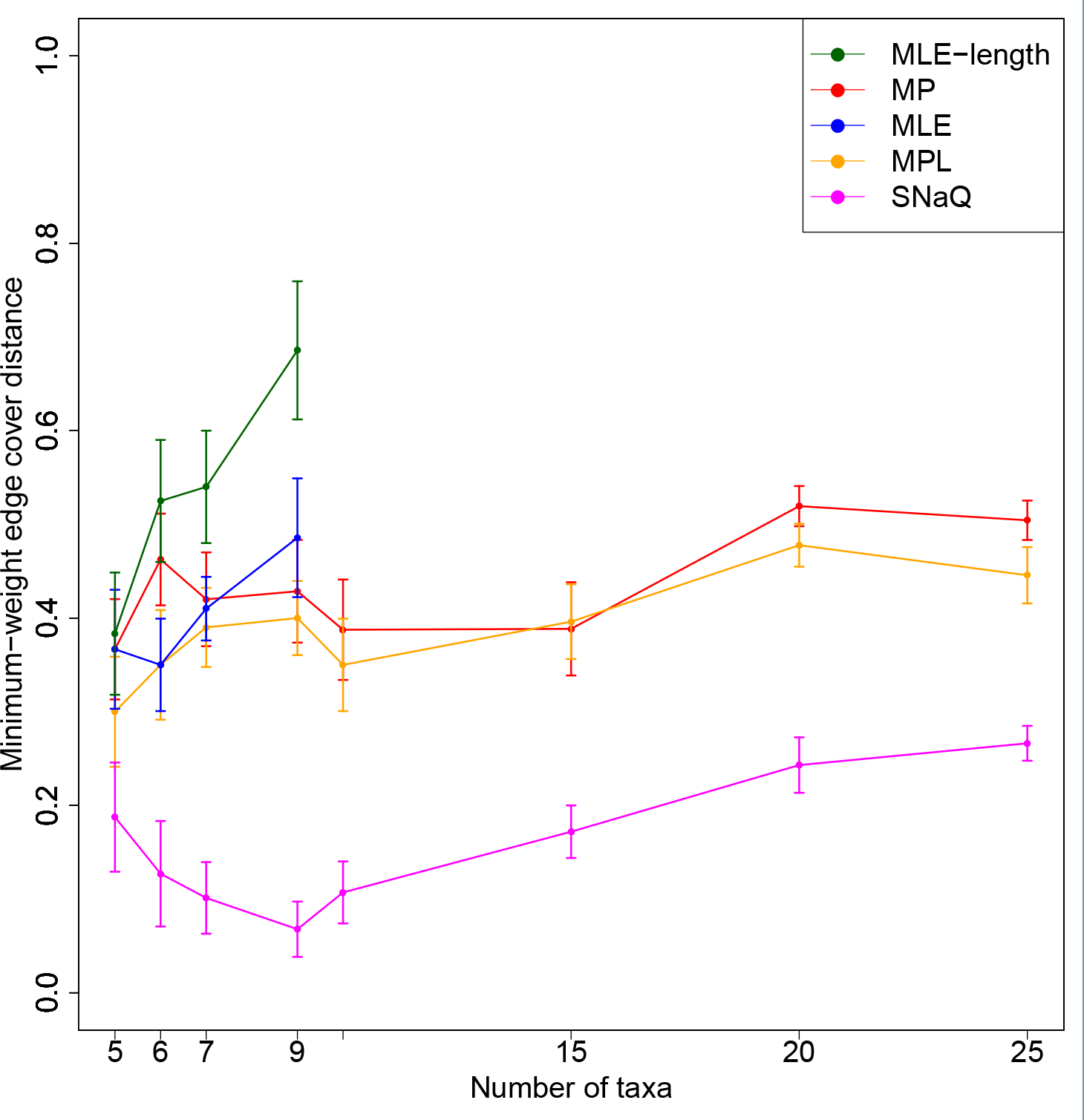
The impact of dataset size on the topological accuracy of multi-locus methods. The model conditions had dataset sizes ranging from 5 to 25 taxa and a mutation rate *θ* of 0.08. Five inferred networks — the phylogenetic networks inferred by a parsimony-based inference method (MP), two full likelihood inference methods (MLE and MLE-length), and two pseudo-likelihood-based inference (MPL and SNaQ) methods — were compared against the model phylogeny in each replicate. The minimum-weight edge cover distance between an inferred network and the model network was used to measure topological accuracy. Average distance and standard error bars are shown (*n* = 20).

**Figure 3.**
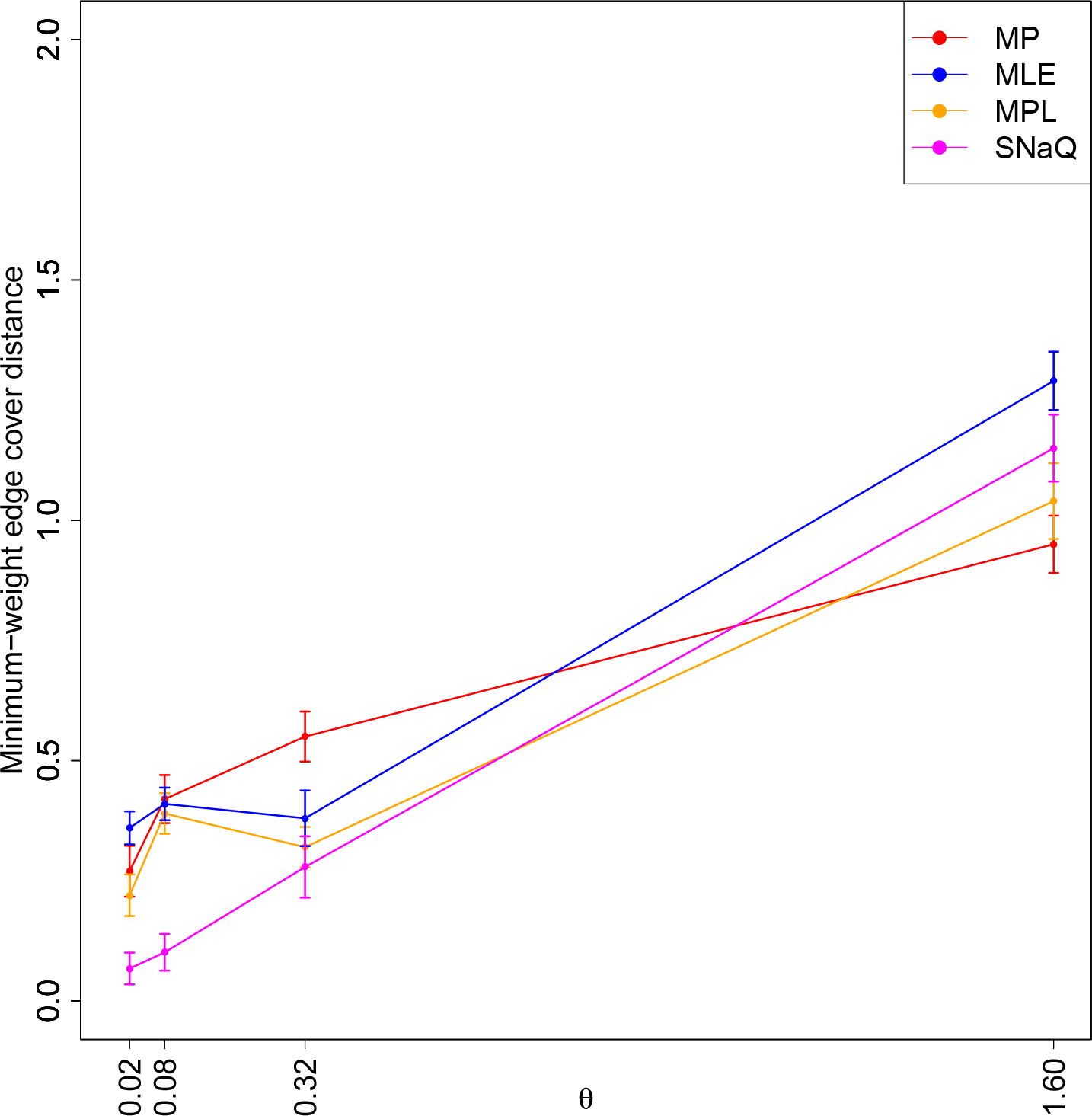
The impact of mutation rate on the topological accuracy of multi-locus inference methods. We assessed the performance of MLE to characterize the full likelihood inference methods since MLE was generally more accurate than MLE-length (Figure 2). The four model conditions had mutation rate *θ* ranging from 0.02 to 1.6 and 7 taxa in each replicate. The minimum-weight edge cover distance between an inferred network and the model network was used to measure topological accuracy. Average distance and standard error bars are shown (*n* = 20).

**Figure 4.**
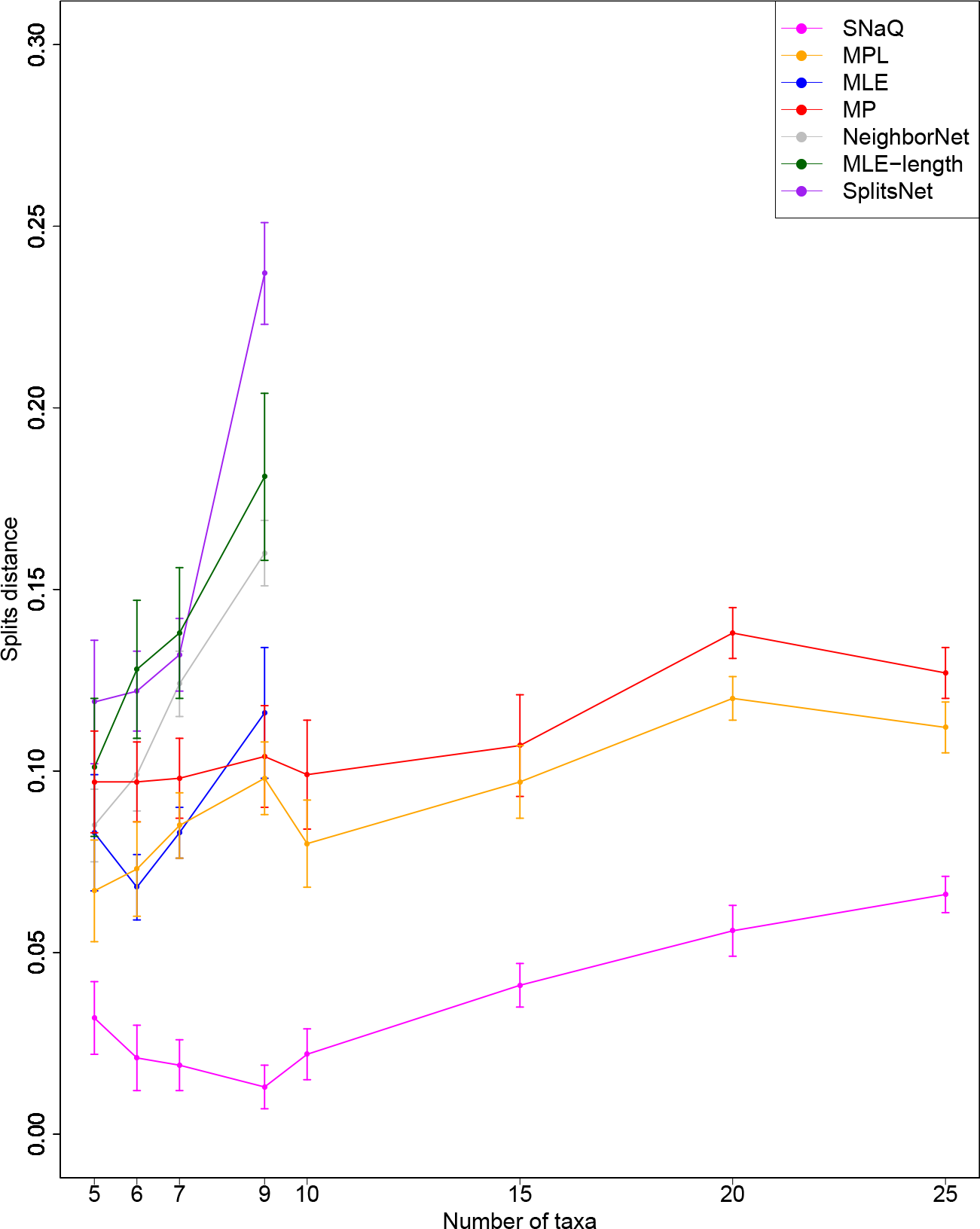
The impact of dataset size on the topological accuracy of the multi-locus inference methods and the concatenated methods. The splits distance between the inferred network and the model network was used to measure topological accuracy. Otherwise, figure layout and description are identical to Figure 2.

Overall, topological accuracy degraded as the number of taxa increased (Figures 2 and 4). Two exceptions to this observation were noted on the smallest datasets in our study: SNaQ’s topological accuracy increased as dataset size increased from five to nine taxa, and a similar general trend was observed with MPL and MP on datasets with between five and ten taxa.

Topological accuracy of the multi-locus methods also degraded as sequence divergence increased due to larger mutation rate *θ* (Figure 3). Compared to the rest of our simulation study, the seven-taxon model condition with mutation rate *θ* = 1.6 was unique for three reasons. First, on this model condition, all methods returned the highest topological error observed in our simulation study. Second, the relative performance of methods on this model condition was unlike the rest of the study: MP, MPL, and SNaQ had comparable topological accuracy (within standard error), and MLE was least accurate. Third, the relative difference in performance was among the smallest observed in our study (e.g., the difference in topological accuracy between the two most accurate methods was smaller on this model condition than any others in our study).

We next examined the impact of gene tree error on the downstream accuracy of the multi-locus inference methods. When gene tree topologies and branch lengths were perfectly accurate, MLE-length was more accurate than all other multi-locus methods, including MLE (Figure 5). In fact, MLE-length inferred topologies that were virtually identical to the model phylogeny. MLE-length’s performance using true gene trees was diametrically opposite to its performance using inferred gene trees, where it was strictly the least accurate among all multi-locus methods. Of the remaining methods, MP was the least accurate overall, and all other methods were intermediate in accuracy with the following exceptions. On the smallest datasets with five taxa, MP and SNaQ were comparable in accuracy. On the largest datasets with nine taxa, MLE-length’s performance advantage over SNaQ diminished and the two had roughly comparable accuracy; furthermore, MPL and MP had comparable accuracy on this model condition. As dataset sizes increased from five to nine taxa, MP and MPL’s accuracy decreased, SNaQ’s accuracy improved, and MLE and MLE-length’s accuracy was largely unchanged. We note that the last observation differs from the experiment using inferred gene trees, whereas the first two observations are largely consistent with the experiment using inferred gene trees. All multi-locus methods were more accurate using true gene trees as input in place of inferred gene trees.

**Figure 5.**
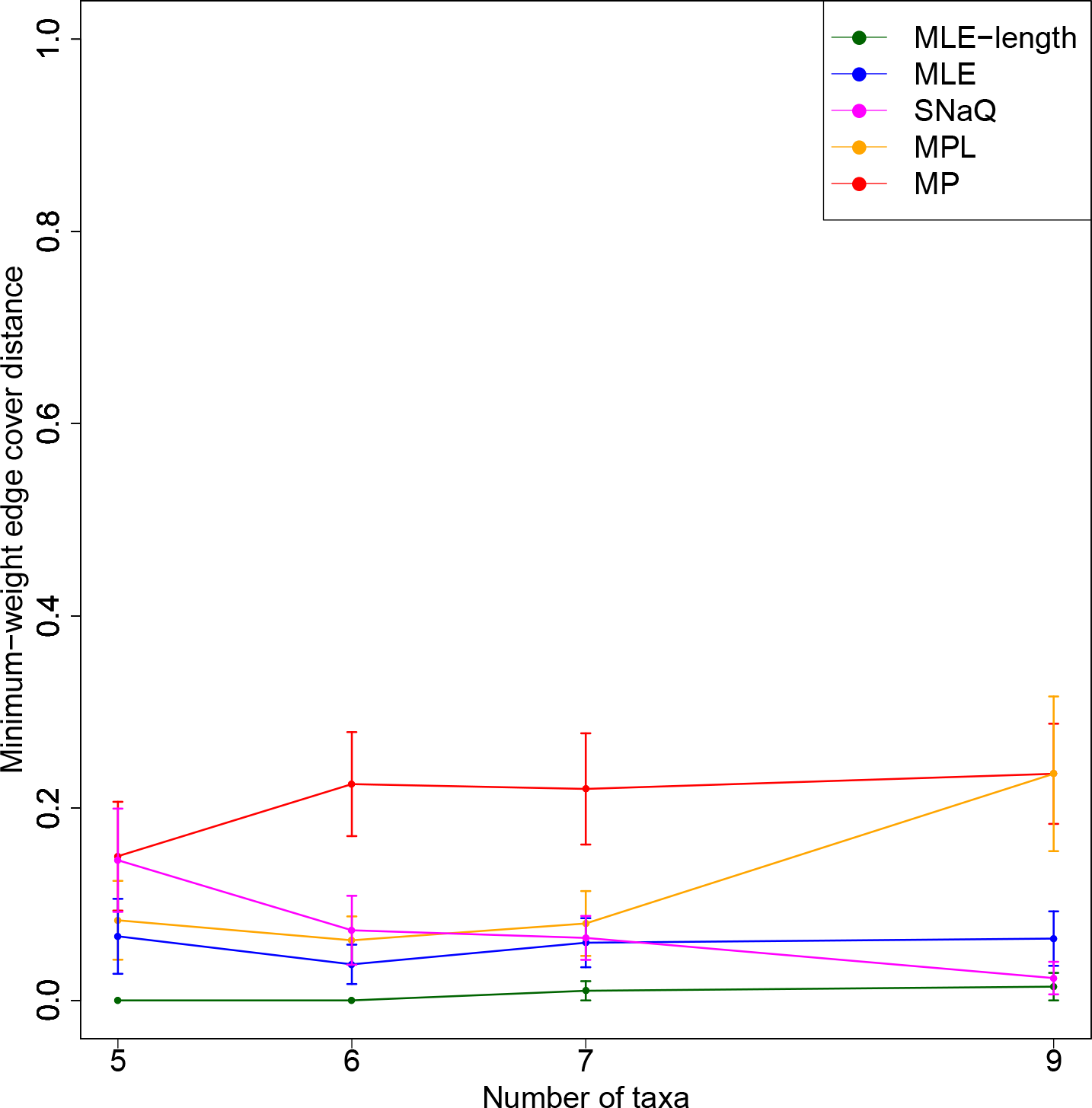
Topological accuracy of multi-locus methods using true gene trees instead of inferred gene trees. The model conditions involved dataset sizes ranging from five to nine taxa. The minimum-weight edge cover distance between an inferred network and the model network was used to measure topological accuracy. Average distances and standard error bars are shown (*n* = 20).

### Performance evaluation on empirical datasets

Our performance study utilized empirical samples from natural populations of *Mus musculus* subspecies and sister species (*M. spretus*, *M. spicilegus*, *M. macedonicus*, and *M. cypriacus*). Prior studies detected gene flow between the *M. musculus* subspecies [42] and between *M. musculus domesticus* and *M. spretus* [43, 44]. We focused our comparison on the most accurate methods from each category of multilocus methods: MLE from the full likelihood methods, SNaQ from the pseudolikelihood-based methods, and MP. We omitted the concatenated methods from our comparison since they were among the least accurate of all methods in our simulation study.

At a coarse level, probabilistic inference using MLE was able to accurately detect gene flow in the empirical datasets. Specifically, the model selection criterion used by MLE consistently chose solutions with gene flow (i.e., phylogenetic networks with one network node) as opposed to solutions without gene flow (i.e., phylogenetic trees). However, as shown in Table 1, all of the methods inferred phylogenies with topologies that differed across replicates (with the exception of the phylogenetic trees inferred by MLE, which were never preferred by the model selection criteria). Greater topological agreement was observed among phylogenies inferred using the same method as opposed to phylogenies inferred using different methods. Furthermore, greater topological agreement was observed when solutions were constrained to have no gene flow, as opposed to solutions involving gene flow. Based on intra-method comparison, the greatest topological agreement was observed among MLE trees, followed by MP trees, SNaQ trees, MP networks, MLE networks, and then SNaQ networks. Based on inter-method comparison, the greatest topological agreement was observed between MP trees and MLE trees, between MP networks and MLE networks, and between MLE networks and SNaQ networks; all other cross-method comparisons involved topological distances that were larger.

**Table 1.** Topological distances between inferred phylogenies in the empirical study. Phylogenies were inferred using the most accurate methods from each category of multi-locus methods: MLE (a full likelihood probabilistic method), MP (a parsimony-based method), and SNaQ (a pseudo-likelihood-based probabilistic method). The top matrix shows the normalized Robinson-Foulds distance between solutions that excluded gene flow (i.e., phylogenetic trees). The bottom matrix shows the normalized minimum weight edge cover distance between solutions that included gene flow (i.e., phylogenetic networks with one network node). Average (standard error) topological distance is shown (*n* = 20). Since the matrices are symmetric, only upper triangular entries are shown.

| Average (SE) topological distance<br>between inferred phylogenetic trees |  |  |  |
| --- | --- | --- | --- |
|  | MLE | MP | SNaQ |
| MLE | 0 (0) | .26 (.01) | .55 (.02) |
| MP |  | .05 (.011) | .79 (.02) |
| SNaQ |  |  | .08 (.009) |

| Average (SE) topological distance<br>between inferred phylogenetic networks |  |  |  |
| --- | --- | --- | --- |
|  | MLE | MP | SNaQ |
| MLE | .183 (.012) | .59 (.03) | 1.16 (.03) |
| MP |  | .156 (.023) | 1.19 (.04) |
| SNaQ |  |  | .295 (.019) |

## Discussion

The probabilistic multi-locus methods were the most accurate methods in our study, but they were also among the most computationally intensive methods in terms of runtime. Given a week of runtime, none of these methods completed analyses of datasets with more than a few dozen taxa. Our finding resolves an open question in the literature: can state-of-the-art phylogenetic network inference methods scale to dataset sizes typically seen in today’s phylogenomic studies? Surprisingly, the answer is no. We were particularly surprised by our finding given the methodological tradeoff made by the pseudo-likelihood-based methods, which use a statistical approximation to full likelihood calculations under the coalescent model to improve scalability. Based on the discussion in [25], we expected that the tradeoff would yield order(s) of magnitude improvements in runtime compared to full likelihood methods, at the cost of reduced topological accuracy. Instead, for datasets with sequence length on the order of 100 kb, the tradeoff only scaled up analyses of around 20 taxa to around 25 taxa. On datasets with more than 30 taxa, the computational requirements of the probabilistic multi-locus methods are projected to be nearing the limits of the most powerful computational clusters available to us. This dataset size is the largest in our study and yet is not considered large in the context of today’s phylogenomic studies. We expect that, like the full likelihood method, the pseudolikelihood-based methods’ memory requirements will grow super-linearly as dataset sizes increase past an inflection point. Finding the inflection point will require additional experiments using larger dataset sizes than those explored in our study. However, we note that the runtime requirements of pseudo-likelihood methods will prove prohibitive on datasets with far more than 25 taxa.

The relative topological accuracy of the methods depended upon whether gene trees were inferred with error-as is the case in virtually all empirical studies-or whether true gene trees were available-a purely theoretical scenario (outside of a few special settings such as experimental evolution in the laboratory). The inferred gene tree error observed in our study (Supplementary Table 2) was comparable to that of other performance studies [45, 46].

When gene trees were inferred, SNaQ was the most accurate method. Our finding overturns a widely held assumption (e.g., [25] asserted that SNaQ was less accurate than MLE and MLE-length). While pseudo-likelihood-based methods were designed to tradeoff inference accuracy for computational efficiency, they are not necessarily less accurate than full likelihood methods. For each replicate, all multi-locus methods used exactly the same set of gene trees as input. We attribute the performance difference to the issue of rooting phylogenies. Prior phylogenetic studies have confirmed the difficulty of accurately rooting phylogenies [47, 48, 49]. Crucially, SNaQ was the only multi-locus method that treated gene trees as unrooted inputs to a quartet-based analysis. In contrast, the other multi-locus methods used rooted gene trees as input and assumed that the root of each gene tree was inferred without error. Intuitively, treating an incorrectly inferred root as correct will propagate error “downstream” during subsequent inference. We note that it is not clear whether using unrooted gene trees is always the best approach. For example, the input for an intermediate approach could utilize a distribution of gene trees for each locus, where the distribution summarized confidence in different rooted versions of unrooted topologies (as well as confidence in different topologies). This approach would strike a balance between two extremes: one where gene tree rooting is completely uncertain (where SNaQ’s approach would be preferred) and the other where there is complete certainty (where the other multi-locus methods’ approach would be reasonable). A Bayesian framework would naturally incorporate gene tree distributions as input.

It is widely assumed in the literature that inference under models incorporating branch length information will be generally more accurate than inference under related models that ignore branch length information [27] (although see the review of Nakhleh [10] for an opposing viewpoint). Our study included probabilistic methods that perform inference under both types of models. In particular, MLE-length and MLE were identical methods with one major exception: the former calculated model likelihood using gene tree topologies and branch lengths, whereas the latter substituted the approach of [23] which calculates model likelihood using only gene tree topologies. When true gene trees were available, MLE-length was the most accurate method. On the other hand, when using inferred gene trees, MLE was more topologically accurate than MLE-length, which we attribute to the difficulty of accurately inferring phylogenetic branch lengths (as noted by [38, 50, 51]). Our findings are consistent with the observations of [14] on a simulation study using a species phylogeny with four taxa (as well as a supplementary set of experiments involving slightly larger species phylogenies); our study more generally shows that the performance comparison holds as dataset size and divergence increases, and further quantifies the impact upon topological accuracy using bipartition-based distance measures. We note that the performance comparison of methods based on their use of branch length information was similar to the comparison of methods that used either rooted or unrooted gene tree inputs. We again observed two extremes: with perfect branch length information (and perfect topological information), MLE-length was strictly more accurate than MLE and the other methods; however, when gene tree branch lengths were inferred with error, treating branch lengths as correct resulted in reduced accuracy compared to methods that ignored branch lengths. Once again, an intermediate approach might offer more flexibility and possible improvements in inference accuracy. One possibility would be inference using a probability distribution over gene tree topologies *and* branch lengths for every locus in a multi-locus dataset. The distributions would reflect confidence in “upstream” inference. Such an approach would likely increase computational requirements even further.

Consistent with other performance studies examining the related problem of scalable phylogenetic tree estimation [17, 52, 53], the parsimony-based multi-locus method were not as accurate as the most accurate probabilistic multi-locus method. The concatenated methods were among the least accurate methods in our study.

Increasing either of the two dimensions of scale-the number of taxa and sequence divergence-generally reduced the topological accuracy of each method (where true gene trees were not available). Both observations are consistent with related studies of phylogenetic tree inference in the presence of gene flow [34, 35]. One contributing factor was inferred gene tree error. Increased sequence divergence due to increasing mutation rates reduced the accuracy of inferred gene trees, which is consistent with theoretical expectations and empirical observations about long branch attraction in other phylogenetic studies [52]. We note that, compared to the effect of increasing the mutation rate, increasing dataset size had relatively little effect upon gene tree inference error but generally increased downstream species phylogeny inference error. The heuristic approaches necessary for analysis of NP-hard optimization problems also contribute to the methods’ scalability; practical issues such as local optima in the search space can pose major challenges to the performance of these heuristics. The relative difference in topological accuracy of gene tree inference compared to species phylogeny inference suggests that the heuristics used for the former performed better than the heuristics used for the latter. One minor exception to these scalability trends was observed. On the smallest datasets, the topological accuracy of SNaQ and MP was either unchanged or improved somewhat as dataset size increased from 5 to 9 taxa. We attribute this finding to long branch attraction. The impact of long branches on parsimony-based phylogenetic inference is well understood [53], and we hypothesize that SNaQ’s use of quartet-based inference may cause it to be more vulnerable to long branch attraction issues compared to the other probabilistic methods in our study.

The findings from our empirical study were consistent with prior studies [44, 42]. Using an information theoretic approach for model selection, MLE consistently inferred historical gene flow between the sampled mouse populations in our study. However, none of the methods were robust to the choice of taxa sampled from the populations under study. This suggests low support which could be due to several causes, including the impact of dataset size and sequence divergence on inference error (consistent with the simulation study) and/or a soft polytomy due to a short branch involving *M. musculus* subspecies (consistent with the consensus phylogeny proposed by Guenet and Bonhomme [54]). Furthermore, the topological distances observed in the empirical study were larger than the topological errors observed in comparable datasets from our simulation study. Even assuming that a species phylogeny inferred on one of the replicates was correct (or close to correct), the topological distances between inferred phylogenies imply that inferences on many of the other replicates would have error comparable to or greater than those observed in the simulation study. One contributing factor is that the empirical datasets may pose a more difficult inference problem since they reflect a broader array of evolutionary processes than those involved in the simulation study. For example, positive selection and recombination have been shown to play significant roles in the evolution of the natural house mouse populations that were sampled in our study [44, 43, 42].

## Conclusions

In this study, we examined the scalability of state-of-the-art phylogenetic network inference methods. We quantified the performance of the methods in terms of computational runtime, main memory usage, and topological accuracy on datasets that varied along two separate dimensions of scale: the number of taxa and sequence divergence.

The methods face tremendous scalability challenges on datasets that are well within the scope of today’s phylogenomic studies. In terms of accuracy, the probabilistic multi-locus methods consistently outperformed the other methods, which is consistent with the state of the art of phylogenetic tree inference. For this reason, we generally recommend using the former-particularly SNaQ, a pseudo-likelihood-based approach-rather than the latter. The latter included concatenated methods-the predominant approach used in today’s phylogenomic studies. More taxa and greater sequence divergence degraded the topological accuracy of all methods. While the probabilistic multi-locus methods retained a performance advantage in terms of topological accuracy, their computational requirements were excessive. On datasets with fewer than 20 taxa and sequence length of 100 kb, the pseudo-likelihood-based multi-locus methods generally completed analysis within a day using around a few GiB of main memory or less; the full likelihood multi-locus method was able to analyze datasets with around 10 taxa within a day and required around 10 GiB of main memory. The computational requirements of the probabilistic multi-locus methods grew rapidly as the number of taxa increased and became prohibitive on datasets with more than 30 taxa. We note that, in a sense, the two dimensions of scale act in opposition: increasing taxon sampling can help reduce evolutionary divergence and inference error, but comes at the cost of increasing the number of taxa which increases runtime requirements and can also increase inference error as well.

We were surprised by several findings which differed from the existing literature. We expected that the pseudo-likelihood-based approaches would scale to much larger datasets than full likelihood methods were capable of analyzing. However, this was clearly not the case. Second, contrary to the assumption of SNaQ’s developers [25], SNaQ’s use of pseudo-likelihood-based calculations did not lead to reduced inference accuracy compared to the full likelihood methods (MLE and MLE-length). In fact, SNaQ was the most accurate method in our study (except in the mostly theoretical situation where true gene trees were available). Third, we came to a conclusion that we feel is understated in the existing literature: when it comes to gene trees, details matter-a lot! The accuracy with which gene trees are rooted appears to impact the downstream accuracy with which species phylogenies are inferred. Furthermore, the accuracy of gene tree branch length inference has a similar effect. The only situation in which MLE-length’s use of gene tree branch length information resulted in more accurate species phylogenies compared to methods that ignored branch length information was when gene tree branch lengths were provided without error. Otherwise, MLE-length was among the least accurate methods in our study.

We highlight several aspects of our study for future work. Most importantly, our study has highlighted the clear need for new phylogenetic inference methods that can cope with the scale of current phylogenomic studies, involving as many as hundreds of genomes; the near future will bring studies that are orders of magnitude larger. We anticipate that our study foreshadows new methodological development on the topic of large-scale phylogenetic network inference. We highlighted one possibility involving methods that performed phylogenetic inference using distributions of gene tree topologies and/or branch lengths as input. We note that such an approach would only magnify the scalability issues that we observed in our study. An expanded empirical study with larger datasets will be possible as future studies follow up on initial reports of gene flow among natural populations (particularly involving different species) and perform additional sequencing. Finally, we propose that the dichotomy between the different categories of methods in our study represents an algorithmic engineering opportunity. By synthesizing these approaches, advantages in one category of methods can help offset disadvantages in the other.

## Methods

### Simulated datasets

*Generation of random model trees using r8s*. Random model trees were generated using r8s version 1.7 [55]. The following script was used to simulate random birth-death model trees for 5, 6, 7, 9, 10, 15, 20, and 25 taxa:

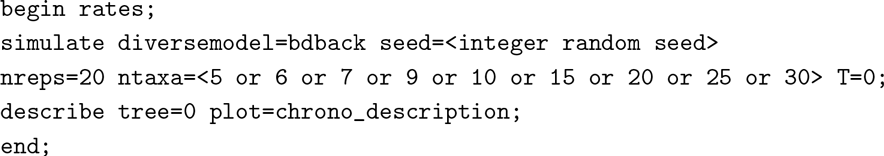

Twenty random model trees (replicates) were generated using r8s. Using a custom script, the branches of each random model tree were scaled by a factor *x* so that the model height phylogeny *h* is 5.

*Generation of random model networks using ms*. We added a single network edge to each random model tree using the following procedure: (1) choose a random time unit t_M_ such that 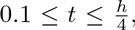, and (2) add unidirectional migration, with a rate of 0.4 which was used in [29], between two taxa or subpopulations such that migration occurs from *t*_*M*_ – 0.1 to *t*_*M*_ + 0.1. A single outgroup was added for each model network at coalescent time 20. We simulated 1000 gene trees for each random model network using ms [56]. The following ms command was used to generate the model network:

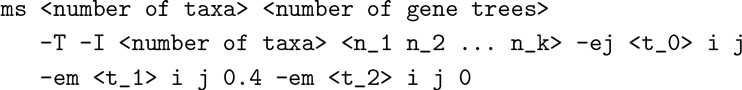

The-T parameter outputs the gene trees that represent the history of the sampled taxa. The-I parameter is followed by *k* that represents the number of subpopulations. The list of integers (*n*_1_ *n*_2_… *n*_*k*_) represents the number of taxa sampled for each subpopulation. We sampled one taxa per subpopulation. The-ej parameter specifies to move all lineages in subpopulation *i* to subpopulation *j* at time *t*_0_. The first-em parameter sets migration at time t_1_ from subpopulation *j* to subpopulation *i* to 0.4. The second-em parameter sets migration at time *t*_2_ from subpopulation *j* to subpopulation *i* to zero.

*Simulation of sequences using seq-gen*. The gene trees output by ms were used as input to seq-gen [57], a sequence evolution program, which can simulate the evolution of sequences according to a finite-sites model. For each local genealogy simulated by ms, we simulated DNA sequence evolution using the Jukes-Cantor mutation model [58]. The total length of the simulated sequences was 100 kb distributed equally across all the local genealogies (100 bp per local genealogy). The following command was used to simulate the evolution of sequences:

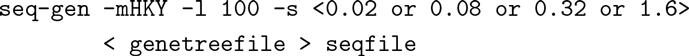

The-mHKY parameter specifies the Jukes-Cantor mutation model. The-s parameter specifies mutation rate *θ* of 0.02, 0.08, 0.32, or 1.6. The-l parameter specifies the length of a sequence in base pairs.

### Phylogenomic inference using inferred gene trees

A single pipeline with two stages was used to infer a species phylogeny.

*Stage one: local gene tree inference using FastTree*. Fast Tree [59, 60] under the Jukes-Cantor model was used to infer the maximum-likelihood unrooted gene tree for each sequence alignment generated by seq-gen. Using a custom script, we converted the branch lengths from expected number of substitutions to coalescent time using equation (3.1) in [61]. The unrooted gene trees were rooted based on the outgroup using PAUP* [62]. Each unrooted gene tree was used as a backbone and the outgroup was added to root each gene tree under the maximum-likelihood criterion. After rooting each inferred gene tree, the outgroup taxon and its pendant edge was pruned.

*Stage two: reconciliation of local gene trees into species phylogeny*. The inferred gene trees were used as input to MLE-length, MLE, MP, MPL, and SNaQ. MLE-length, MLE, MP, and MPL are implemented as part of the PhyloNet [22] package. The following is a sample NEXUS script file that was used to execute the PhyloNet commands:

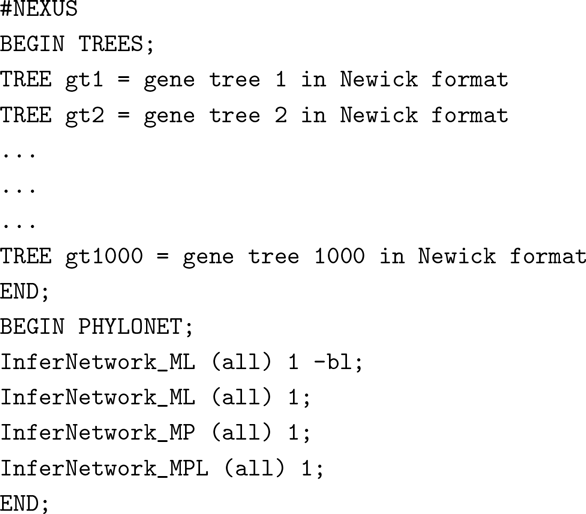

The commands located in the `TREES` block contain the inferred gene trees. The commands located in the `PHYLONET` block contain the inference methods and parameters used to infer a species network. The InferNetwork_ML command infers a species network with one reticulation node using maximum likelihood. The-bl parameter specifies the use of branch lengths of gene trees in the inference. In the absence of-bl, only the topologies of gene trees are used in the inference. The In-ferNetwork_MP command infers a species network with one reticulation node using a parsimony-based method under the MDC criterion. The InferNetwork_MPL command infers a species network with one reticulation node using maximum pseudolikelihood.

The following is a sample script used to execute the SNaQ commands:

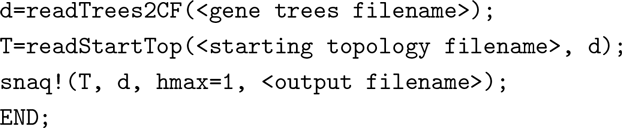

The gene trees are summarized as quartet concordance factors using the readTrees2CF function. The readStartTop reads the tree used as a starting point for the search. The starting tree was estimated using the MDC criterion. The snaq! command estimates a network using the input quartet concordance factors *T* and starting from tree *d. hmax* specifies the number of reticulation nodes.

### Phylogenomic inference using true gene trees

The true gene trees generated by ms were used as input to the following phylogenomic inference methods: MLE-length, MLE, MP, MPL, and SNaQ.

### Concatenated analysis

For the concatenated analyses, we inferred species networks using two distance-based methods implemented in the phangorn software package [63]: (1) NeighborNet [11], a clustering method that extends the neighbor-joining algorithm, and (2) the least squares method of Schliep [12]. Throughout this manuscript, we refer to the latter as SplitsNet, since it infers a split graph. The Hamming distance matrix was used as input to the distance-based concatenated methods.

### Measuring accuracy

Accuracy was computed by comparing the inferred phylogeny to the model phylogeny using minimum-weight edge cover [64]. This measure compares the similarity between the set of trees induced by the inferred and model networks using Robinson-Foulds (RF) distance [65], and then identifies the minimum sum of weights in a bipartite graph where a weight is the RF distance between a tree induced in the inferred network and a tree induced in the model network. The RF distance counts the number of false positive bipartitions (bipartitions found only in the inferred network) and false negative bipartitions (bipartitions found only in the model network). A tripartition-based measure, which finds the proportion of tripartitions that are not shared between two networks, was also used to compute the distance between an inferred and model rooted networks [64]. Finally, the splits distance, which identifies the proportion of bipartitions found in the inferred species network but not in true species network and proportion of bipartitions found in true species network but not in inferred species network, was used. The second evaluation criteria used was the computational requirements of the inference methods, which was measured in terms of running CPU time and memory usage.

### Empirical datasets

We used genomic sequence data sampled from natural mouse populations. A recent study has highlighted historical gene flow between some of the populations in our study [3]. The samples were collected in previous studies [3, 42, 66, 43, 67, 68]. The collected sample information contained 100 haploid mouse genomes that are either wild or wild-derived samples. The procedure that was used to generate the sequence data is described in [3]. The sequences were filtered to 414,376 SNPs that were genotyped across all samples.

Datasets were constructed from the empirical samples using the following sampling procedure. For each dataset, we randomly selected a sample from each of the following mouse species or subspecies: *Mus musculus domesticus*, *M. musculus musculus*, *M. musculus castaneus*, *M. spretus*, *M. spicilegus*, *M. macedonicus*, and *M. cypriacus*. The sampling was repeated twenty times to obtain twenty datasets.

We estimated recombination-free intervals for use as the input loci to the phylogenetic network inference methods. This required inferring recombination breakpoints. We obtained breakpoints using RecHMM, a hidden Markov model-based method [69], resulting in 3013 recombination-free genomic regions. FastTree using Generalized Time-Reversible model [70] was used to infer the gene tree for each recombination-free genomic region resulting in 3013 gene trees. We used rat (the rn5 assembly downloaded from the UCSC Genome Browser [71]) as an outgroup to root each gene tree generated by FastTree.

MLE, MPL, MP, and SNaQ were used to infer species networks with zero or one reticulation nodes. For inferred networks with zero reticulation nodes, Robinson-Foulds distance was used to measure the topological distance between all inferred network replicates. For inferred networks with one reticulation node, the minimum weight edge cover distance was used to compute the topological distance between all inferred network replicates. The tripartition measure was also used to compute the distance between all inferred network replicates with one reticulation node for the MLE, MPL, and MP methods. For the concatenated methods (Neighbor-Net and SplitsNet), we used the splits distance to measure the topological distance between all inferred network replicates.

### Availability of software and data

All software implementations and datasets are publicly available under open license. More information and download URLs can be found at https://gitlab.msu.edu/liulab/network-inference-scalability-study/wikis/home.

### Competing interests

The authors declare that they have no competing interests.

## List of abbreviations

ILS: incomplete lineage sorting
MP: maximum parsimony
MDC: minimize deep coalescence
MLE: maximum likelihood estimation
MPL: maximum pseudo-likelihood
SNaQ: Species Networks applying Quartets
HMM: hidden Markov model
RF distance: Robinson-Foulds distance

## Author’s contributions

Conceived and designed the experiments: HAH KJL. Implemented software tools: HAH. Performed the experiments: HAH. Analyzed the data: HAH KJL. Wrote the paper: HAH KJL. All authors read and approved the final manuscript.

## Acknowledgments

This work is supported by startup funds from Michigan State University to KJL.

